# Supershoes or Superhumans? A controlled analysis of sex-specific road-running performance evolution in the era of advanced footwear technology

**DOI:** 10.64898/2026.07.03.736031

**Authors:** Leonie Blattmann, Daniel Hamacher, Ross Tucker, Laura Healey, Joel Mason

## Abstract

Advanced footwear technology (AFT) improves running economy and is widely credited with recent performance improvements in road running, with observational analyses consistently demonstrating that women have improved more than men. In contrast, laboratory studies generally report similar running economy responses to AFT in both sexes, leaving the observational-experimental divergence unresolved. A fundamental limitation of observational work is the absence of a control condition, making it impossible to separate AFT-related gains from concurrent performance trends. We addressed this by comparing performances between pre-AFT (2009-2015) and AFT (2017-2024) eras across road-running events (10km, half marathon, marathon) and throwing events (shot put, discus, javelin, hammer), the latter serving as an active control condition subject to the same broad athletic trends but unaffected by footwear technology. The top 50 performances per event, era, and sex were converted to World Athletics points and analysed using linear mixed-effects models. Performances improved significantly between eras (β = 0.294, p < .001), with gains substantially larger in road running than in throwing (β = 0.731, p < .001). The sex-specific pattern of improvement also differed between event categories (era × event type × sex interaction, β = 0.786, p < .001): road-running improvements were greater in women than men (4.51% vs 2.62%), whereas throwing improvements did not differ by sex (0.93% vs 1.14%). These findings suggest that AFT benefits women more than men in competition, whether through a greater physiological response or more effective translation of economy gains to race performance, and suggest current laboratory protocols may be insufficiently sensitive to detect potential sex-specific effects.

## Introduction

Following a period of relative performance stability across track and field events (Berthelot et al., 2010, 2015; Ganse & Degens, 2021), recent analyses indicate that elite running performances have exceeded expected trajectories (Needles & Grabowski, 2024; Willwacher et al. 2024). The emergence of advanced footwear technology (AFT) in 2016 is widely considered a key driver of these latest evolutions in performance. AFT combines highly compliant and resilient foams with rigid components (Frederick, 2022) to reduce the metabolic cost of running at a given speed (Hoogkamer et al., 2018; Hunter et al., 2019) and ultimately improve running performance.

Ecological evidence is consistent with a substantial effect of AFT in road running. Analyses of annual top lists have documented performance improvements of 1-3% in distances ranging from 10km to marathon in the AFT era (Bermon et al., 2021; Senefeld et al., 2021; Rodrigo-Carranza et al., 2021; Langley & Langley, 2024; Langley et al., 2023). Beyond these general performance improvements, observational analyses consistently reveal that women have experienced greater performance improvements than men since the release of AFT (Mason et al., 2024; Senefeld et al., 2021; Bermon et al., 2021; Willwacher et al., 2024). Women’s road-running world records have improved by an average of 3.7% since 2016 compared to 1.5% in equivalent men’s events (Mason et al., 2024), annual top 20 and top 100 progressions show similar asymmetries (Bermon et al., 2021), and within-athlete comparisons of 239 marathoners switching from conventional to AFT footwear show women gaining roughly twice as much as men (1.6% vs 0.8%; Senefeld et al., 2021). This marks a notable departure from a sex gap in running that had been stable since the 1990s (Cheuvront et al., 2005; Millard-Stafford, Swanson & Wittbrodt, 2018; Thibault et al. 2010).

These consistent observational findings contrast with experimental evidence examining running economy, the key mechanism through which AFT improves running performance (Hoogkamer et al., 2018). In laboratory comparisons of AFT versus conventional footwear, direct intrastudy male-female comparisons have consistently reported no sex differences in the metabolic responses to AFT at matched speeds in highly trained runners (Barnes & Kilding, 2019; Seglina et al., 2026) and in amateur triathletes (Riedl et al., 2024). Of these studies, only Seglina and colleagues specifically designed their work to detect sex differences. Although women remain underrepresented in experimental AFT datasets (Mason et al., 2024), syntheses of currently available evidence conclude that sex is not a predictor of running economy responses to AFT (Stephen et al., 2025; Burns & Joubert, 2024). Against this consensus, Matties and Rowley (2025) recently reported that female collegiate runners experienced a significantly greater energetic cost benefit from AFT (∼4.7%) than male runners (∼2.8%) when testing speed was controlled. This remains the only experimental evidence of sex-specific responses to AFT.

The prevailing divergence between observational and experimental evidence may emerge because experiments and observational analyses have been asking different questions. Experiments ask how AFT alters physiology, while observational analyses ask whether AFT can help explain what we see in competition, and whether it can help explain why women have gained more than men. However, observational analyses cannot rule out the possibility that factors beyond footwear have also contributed to recent performance changes. This possibility is highlighted by a recent preprint analysis arguing that longer-term changes in opportunity, participation, professionalisation and competitive depth explain women’s disproportionate running gains instead of AFT (Jucker, 2026). Yet both sides of this debate rest on the same uncontrolled foundation: without a non-footwear-exposed comparator condition, they cannot determine whether the recent sex-specific evolution of road-running performance reflects footwear, broader changes in women’s sporting performances, or both.

To address this challenge, we examined throwing events as an active control group. Contested within the same elite athletics ecosystem as road running, throwing is subject to the same era-level forces shaping the sport, including evolving training methods, professionalisation, doping, participation trends and broader societal change (Haake, James & Foster, 2014), but remains unaffected by the recent advances in footwear technology, making it well suited to isolating the contribution of AFT from concurrent general influences on performance. We hypothesised that: (1) both road running and throwing events have improved between eras, with road running demonstrating greater gains than throwing regardless of sex, (2) women have improved more than men in both event categories, and (3) the magnitude of sex-specific improvements is greater in running than in throwing events, suggesting AFT-specific benefits that are additive to broader athletic trends.

## Methods

### Experimental Approach and Data Selection

To address the study aim, seven track and field events were analysed and categorised into two categories: road running events with AFT available (10km, half marathon, marathon) and throwing events without AFT available (shot put, hammer throw, discus, javelin). Throwing events served as an active control group to control for time effects. They were selected because they are contested within the same elite athletics ecosystem as road running and are subject to the same broad era-level forces shaping the sport but remain unaffected by advances in shoe technology. Performance data from 2009 to 2024 were divided into two eras based on the introduction of AFT in 2016 for Road Running. The Pre-AFT era ranged from 2009 to 2015, while the AFT era covered 2017 to 2024. The year 2016 served as the transition point and was excluded from the analysis as it represents the initial prototype and commercial release period when AFT was available but not yet widely adopted by elite athletes. Additionally, in line with previous analyses of annual performances (Mason et al., 2023), 2020 was excluded from the analysis due to the COVID-19 pandemic, which substantially limited training and competition opportunities. Therefore, both eras consisted of seven years.

Performance data were extracted from the World Athletics All-Time Top Lists (World Athletics, 2025). For each event, the 50 best performances in each era were identified separately for male and female athletes, representing the elite level of performance. Only officially recognised performances were included, with one performance per athlete per event and era selected. If an athlete appeared in both eras, their best performance in each era was included for analysis.

### Data Processing and Statistical Analysis

To enable standardised statistical comparison across different event types with varying units of measurement, all performances were converted to World Athletics Points (WA Points). The World Athletics Points system provides a standardised approach to compare performances across different track and field events by converting results into point values that represent equivalent performance levels (World Athletics, 2017). Since the World Athletics scoring tables were recalibrated in 2025 to account for AFT-related performance benefits suggested in previous research, the scoring tables from 2017 were used for the current analysis to ensure that performance points were assigned independently of footwear technology (World Athletics, 2017). For each event, raw performance data was retained for event-specific further analyses and data visualisation.

The variables were transformed prior to statistical analysis. To account for systematic differences between events, WA Points were z-standardised within each event, which ensured that means and standard deviations were comparable across events. The z-standardized WA Points were then analysed using a linear mixed-effects model. Heterogeneity in the nested data structure was addressed by including multiple random intercepts: because performance levels may differ across ranks even within a given event, a random intercept for each rank nested within each event was specified. In addition, since some athletes participated in multiple events, further random intercept for each athlete was included to account for the dependence introduced by repeated observations of the same individual.

The specification of the mixed-effects model was guided by the study hypotheses. To address Hypothesis 1, we examined (a) whether WA points differ between eras (pre-AFT vs. AFT era) and (b) whether the improvement across eras is greater in AFT-affected events (running) compared to events not affected by AFT (throwing). Therefore, era and the interaction between era (pre-AFT vs. AFT) and event type (running vs. throwing) were included as predictors. In accordance with the Principle of Marginality, the main effect of event type was also included in the model. Hypothesis 2 posits that women may have improved their performance more than men across eras; therefore, factor sex (men/women) was included as an additional predictor. Hypothesis 3 examines whether women benefited even more from the AFT effect than men. To evaluate this, we tested the three-way interaction between era × event type × sex. In line with the Principle of Marginality, all lower-order interaction terms were also included in the model.

Because specifying random effects for these predictors would lead to overparameterisation (rendering the model non-estimable), all predictors were modelled as fixed effects. This leads to the following model equation for the mixed-effects model. For observation *i* in event *k*, rank group ℓ, and athlete *m*, the mixed-effects model is specified as follows:

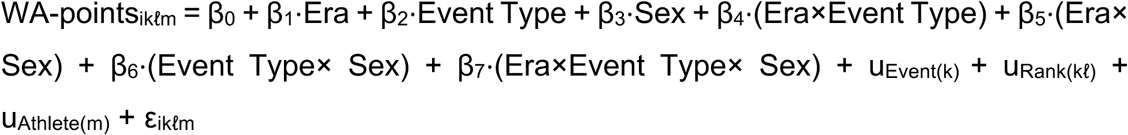

Here, u specifies the random intercepts, and *ε* represents the level-1 residual.

In a secondary analysis, the effect of era on WA points was tested separately for each sex and each event using linear mixed-effects models. In these models, era was the sole fixed effect predictor, and random intercepts were included for athletes.

All model parameters were estimated using restricted maximum likelihood (REML). P-values were obtained from t-tests based on Satterthwaite’s approximation of the degrees of freedom. Effects were considered statistically significant when p < 0.05 (α = 5%).

To provide historical context beyond the era-based comparisons, we also performed an exploratory descriptive churn analysis of all-time lists. For each road-running and throwing event, we identified how many performances achieved during the pre-AFT era remained in the current top 50 all-time rankings as of 31 December 2025. Similarly to the main analysis, performance data were extracted from the World Athletics All-Time Top Lists (World Athletics, 2025).

## Results

### Grouped Analysis: Road-Running vs Throwing Evolutions in the AFT era

According to the mixed-effects model (Table 1, Figure 1), a significant main effect of era was observed (β = 0.294, p < .001), indicating an improvement in throwing, the reference category, between the pre-AFT and AFT eras. The era × sex interaction was not significant (β = –0.019, p = .701), indicating that the era-related improvement in throwing events did not differ between men and women. By contrast, road running improved substantially more than throwing (era × event type, β = 0.731, p < .001), and this running advantage was itself greater in women than in men (era × event type × sex, β = 0.786, p < .001).

**Figure 1:**
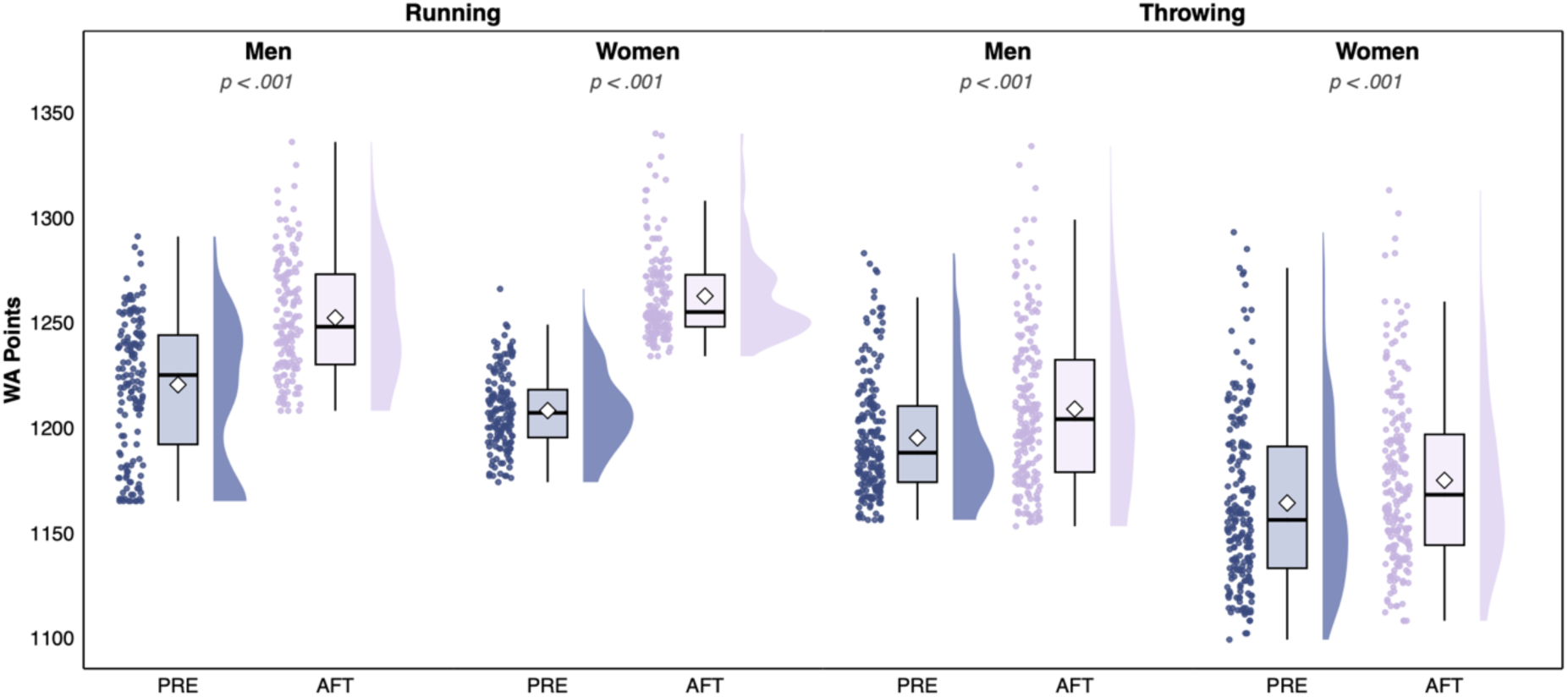
World Athletics (WA) points by event group, era (pre-AFT, 2009-2015; AFT, 2017-2024), and sex. Each group pools the top 50 performances from its constituent events (road-running: 3 events; throwing: 4 events), so each box summarises 150 and 200 performances respectively. Diamonds indicate means, boxes interquartile ranges; each dot is one performance.

**Table 1:** Linear mixed-effects model testing the effects of era, event type, sex, and their interactions on z-standardised World Athletics (WA) points. Model parameters were estimated using restricted maximum likelihood. Degrees of freedom were approximated using Satterthwaite’s method.

| Fixed effect | Estimate | SE | df | t-value | p-value |
| --- | --- | --- | --- | --- | --- |
| **Intercept** | 0.289 | 0.067 | 348.4 | 4.307 | <.001 |
| **Era (factor)**<br>(ref. group: "pre-AFT") | 0.294 | 0.036 | 1109.0 | 8.240 | <.001 |
| **Event type (factor)**<br>(ref. group: "throwing") | -0.741 | 0.101 | 356.0 | -7.361 | <.001 |
| **Sex (factor)**<br>(ref. group: "men") | -0.769 | 0.036 | 1109.0 | -21.563 | <.001 |
| **Era x event type** | 0.731 | 0.055 | 1111.5 | 13.315 | <.001 |
| **Era x sex** | -0.019 | 0.050 | 1109.0 | -0.385 | .701 |
| **Event type x sex** | 0.294 | 0.054 | 1109.0 | 5.393 | <.001 |
| **Era x event type x sex** | 0.786 | 0.077 | 1109.0 | 10.197 | <.001 |

| Random effects | Variance |
| --- | --- |
| Athlete | <0.001 |
| Rank (nested within event) | 0.602 |
| Event | <0.001 |
| Residual | 0.127 |

**Notes**
- SE = standard error - df = degrees of freedom - ref. group = reference group of a dichotomous predictor (baseline category)

In descriptive terms, performances in road-running events improved by 4.51% in women and 2.62% in men, corresponding to mean gains of 44.3 and 18.7 World Athletics points respectively. Throwing improvements were smaller and similar between the sexes (women’s improvement between eras: 0.93%; men’s: 1.14%).

### Individual Event Analysis in the AFT era

Secondary event-specific analyses broadly supported the grouped findings. In road running, all six of the event-sex comparisons improved significantly between eras, both when assessed using raw performances and using World Athletics Points (Table 2, Figures 2-3). Men’s road running time improvements between eras ranged from .90% in the half marathon to 1.75% in the 10km, whereas women’s time progressions between eras ranged from 3.04% (marathon) to 3.52% (10km) (Table 2). In contrast, throwing events showed smaller and less consistent changes. Of the eight event-sex comparisons, four improved significantly (men’s shotput, discus, and javelin; women’s discus), three showed no significant change (women’s shotput, javelin, and hammer), and one showed a significant decrement (men’s hammer; Table 2). Notably, men’s shotput showed the largest throwing improvement at 2.72%, exceeding the improvements observed in men’s marathon and half marathon.

**Figure 2:**
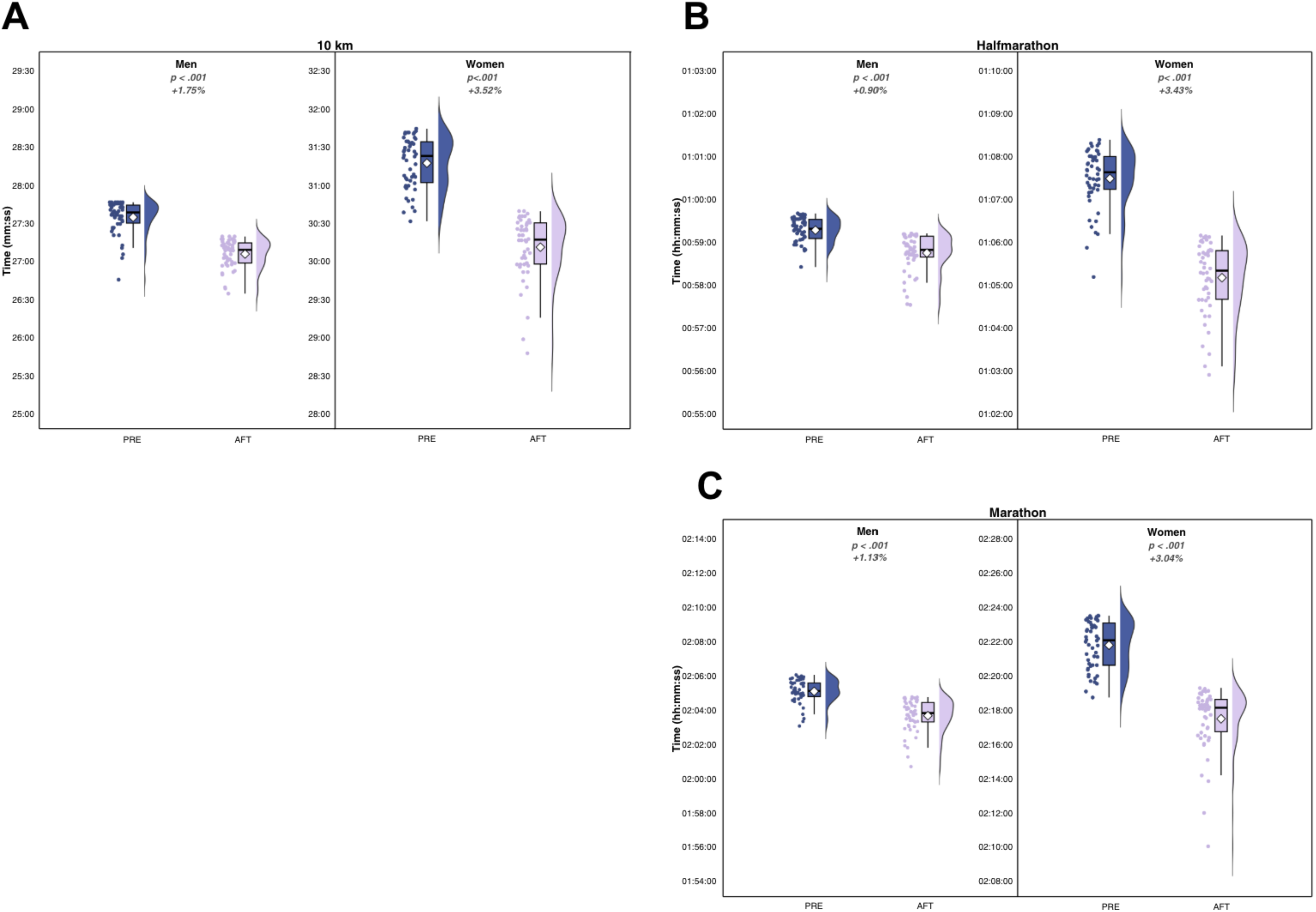
Raw performances for the top 50 athletes in the pre-AFT era (2009-2015) and AFT era (2017-2024) for the. **A:** 10km, **B:** half marathon, and **C:** marathon, shown separately for men and women. Diamonds indicate means, boxes indicate interquartile ranges, and each dot represents one performance.

**Figure 3:**
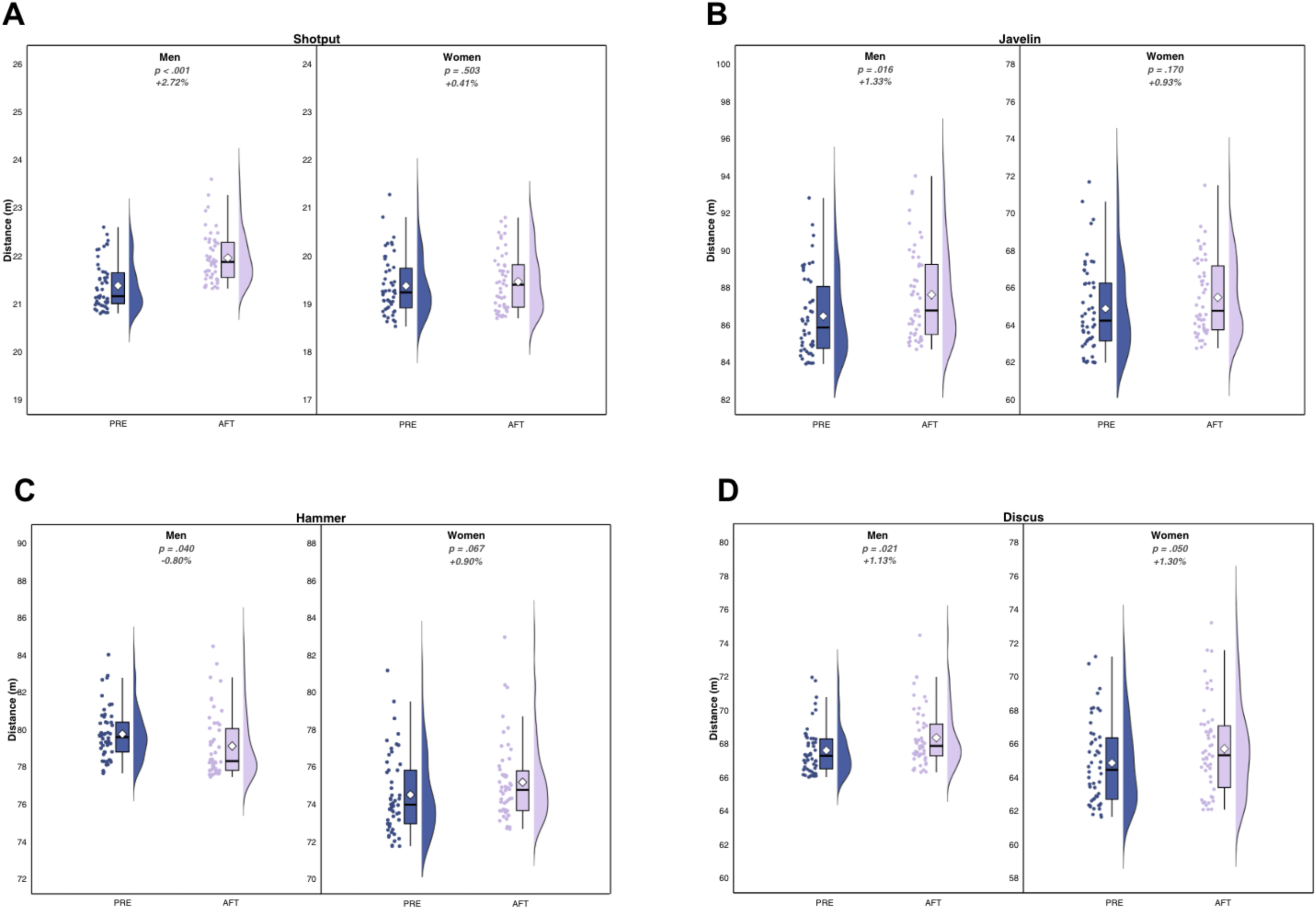
Raw performances for the top 50 athletes in the pre-AFT era (2009-2015) and AFT era (2017-2024) for the control events, **A:** Shot put, **B:** Javelin, **C:** Hammer and **D:** Discus, shown separately for men and women. Diamonds indicate means, boxes indicate interquartile ranges, and each dot represents a single performance.

**Table 2:**
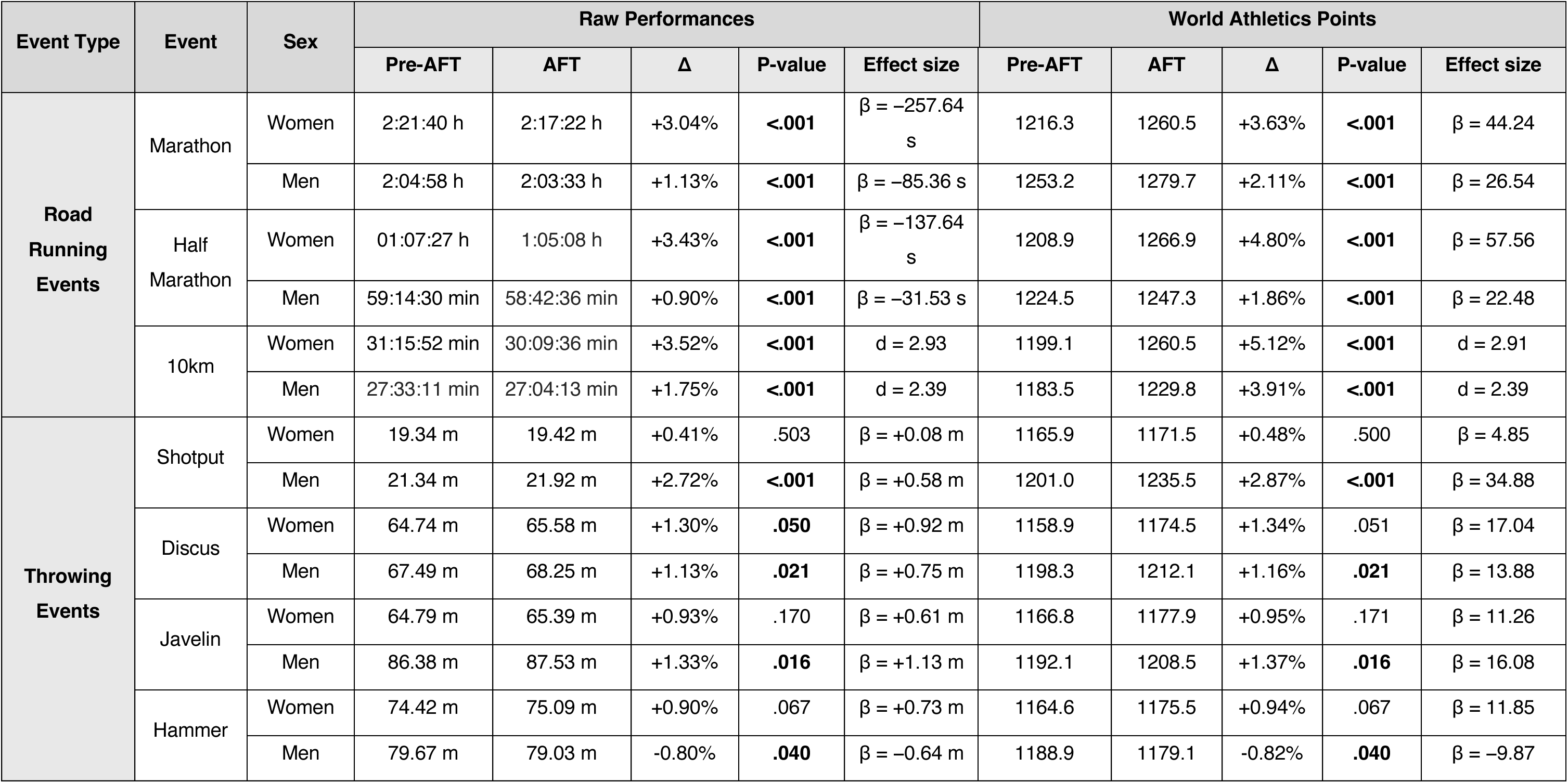
Mean raw performances and World Athletics points for the top 50 performances in each event during the pre-AFT era (2009-2015) and AFT era (2017-2024), shown separately for men and women. Effect sizes are reported as slopes, except where no athletes appeared in both eras, in which case Cohen’s *d* is reported.

### Exploratory Churn Analysis: Composition of current all-time performances

As an additional exploratory analysis to provide descriptive historical context for the era-based comparisons, we quantified the survival of pre-AFT performances in current top 50 all-time lists for each road-running and throwing discipline. Across road-running events, pre-AFT performances were markedly less likely to remain in contemporary all-time rankings than in throwing events, indicating substantially greater turnover in the AFT era. Survival of pre-AFT performances in the AFT era for throwing events ranged from 56%-86% for men and 50%-92% for women. In contrast, for road-running, survival of pre-AFT performances in the all-time lists was at 8.9% (10km), 10.7% (half marathon) and 12.0% (marathon) for men, and 0% (10km), 2% (half marathon) and 2% (marathon) for women (Figure 4).

**Figure 4:**
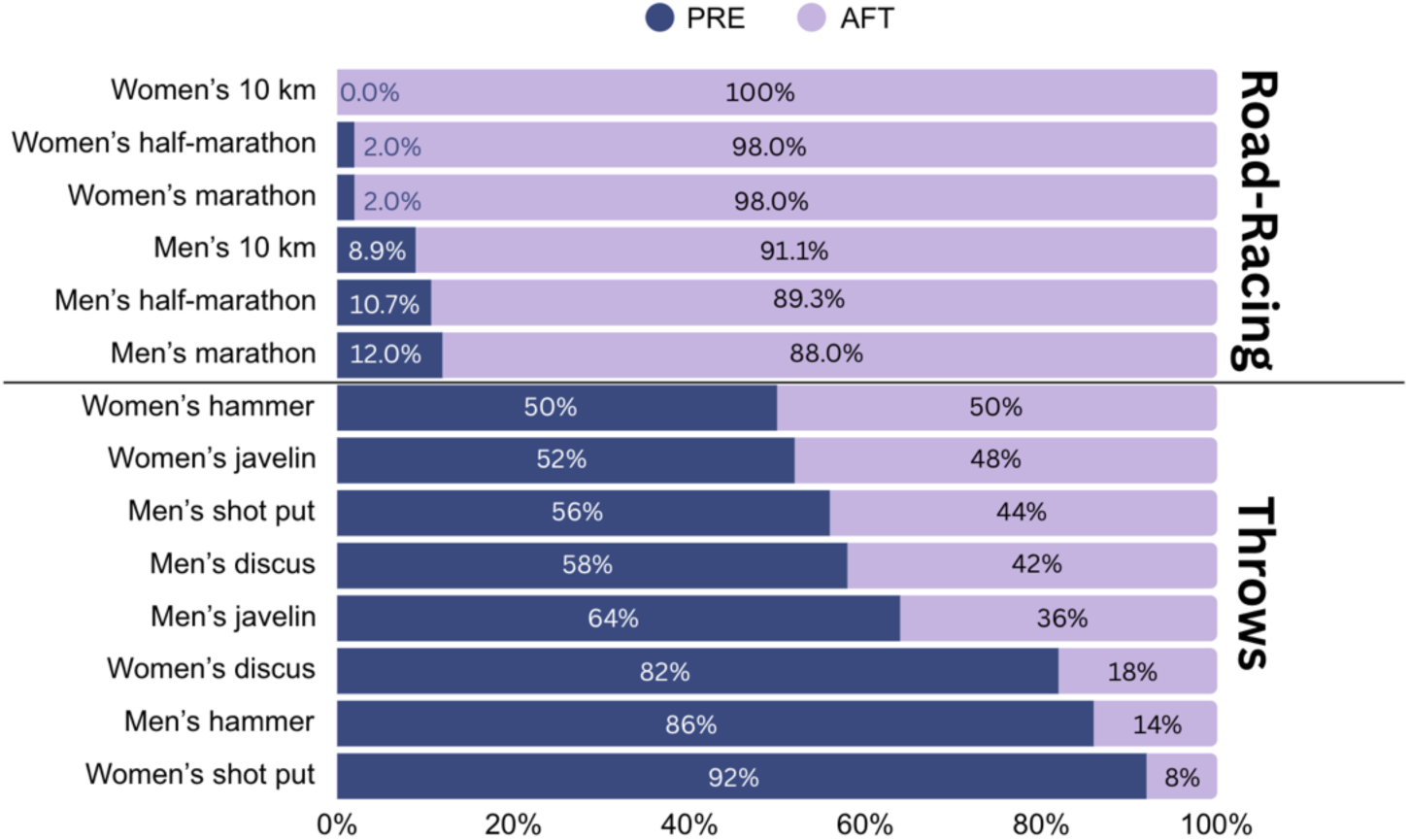
Composition of current top 50 all-time performances by era, shown separately for each event and sex. Dark bars (PRE) represent the proportion of current top 50 performances achieved during the pre-AFT era (any time before 2016), and light bars (AFT) represent the proportion achieved during the AFT era (2017-current). Road-racing events show almost complete displacement of pre-AFT performances from contemporary rankings, particularly in women’s events. In contrast, throwing events (bottom) retain a substantial proportion of pre-AFT performances, consistent with substantially lower era-related turnover. Where performances were tied at the 50th position, all tied performances were included in percentage calculations.

## Discussion

Using the World Athletics points system as a common currency across disciplines and leveraging throwing events as an active control condition, we investigated sex– and discipline-specific performance progressions across the pre-AFT and AFT eras. Performances improved across both road-running and throwing events in the AFT era, confirming an upward trend in elite athletics over this period. However, road-running improved substantially more than throwing events between eras (3.58% vs. 1.04%), and women improved more than men in road-running events (4.51% vs. 2.62%) despite similar improvements between the sexes in throwing events (women: 0.93%, men: 1.14%). Additionally, our churn analysis provided historical context by showing that pre-AFT performances were far less likely to persist in contemporary all-time road-running lists than in throwing events. If broader improvements in women’s sport beyond footwear were responsible for the recent sex-specific gains in road running, we would have expected a comparable female advantage to emerge in the throwing events, which share the same general era-level influences. We did not observe this. In providing a non-footwear-exposed active control for the first time, our findings constitute the most compelling ecological evidence to date that AFT confers greater competitive benefit to women than men.

These findings align with prior observational analyses showing disproportionate gains in women’s road running since the introduction of AFT (Bermon et al., 2021; Senefeld et al., 2021; Goss et al., 2022; Willwacher et al., 2024; Mason et al. 2024), while extending that literature by providing a more controlled comparison against disciplines not exposed to the same technological shift. In doing so, these findings challenge the prevailing interpretation from experimental studies that men and women respond similarly to AFT (Barnes & Kilding, 2019; Seglina et al., 2026; Riedl et al., 2024; Stephen et al. 2025) and support the existence of sex-specific AFT advantages (Mason et al., 2024; Matties & Rowley, 2025) that remain largely undetected in conventional lab-based protocols.

### Reconciling observational and experimental contradictions

Two broad explanations may account for this apparent contradiction between experimental physiological evidence and ecological racing evidence. First, most current laboratory protocols may not be sufficiently sensitive to detect sex-specific responses to AFT, whether those differences arise in the acute physiological response itself, in the translation of that running economy response to race-day performance, or in both. Second, the larger improvements observed in women’s road-running performance may reflect factors beyond footwear technology that have disproportionately influenced female endurance performance in recent years. These explanations are not mutually exclusive, but the absence of a comparable sex-specific pattern in throwing events supports the notion that footwear is, at least to some degree, responsible for the disproportionately large performance improvements in women’s road running events. We therefore explore why standard laboratory protocols may fail to capture the observational results here, before turning to the alternative explanations that our design cannot fully exclude.

### Experimental barriers to detecting potential sex differences in AFT response

Controlled laboratory studies have been instrumental in identifying the benefits of AFT, particularly in establishing improved running economy as the primary physiological pathway underpinning footwear-related performance advantages (Hoogkamer et al., 2016; Hoogkamer et al., 2018). However, the same protocols that made this work possible, typically short, steady-state trials performed in a fresh state, are also the source of their principal limitation. While offering the experimental control necessary for precise measurement, they do not fully capture the physiological and performance demands of road racing lasting from 26 and a half minutes to over two hours.

One pathway by which the laboratory-to-race divergence may arise is that similar metabolic benefits do not necessarily translate into similar performance benefits (Kipp et al., 2019). Running economy can predict performance changes in response to footwear modifications (Hoogkamer et al., 2016), though this foundational work was conducted exclusively in men prior to the introduction of AFT. Notably, competitive advantages from AFT have been demonstrated to exceed what running economy improvements alone would predict. Hébert-Losier et al. (2022) found that the Nike Vaporfly 4% improved running economy by 1.0-1.7% in laboratory conditions, yet 3km time trial performance improved by 1.8%. Similarly, Black et al. (2022) observed that highly cushioned running shoes improved incremental treadmill test performance by 5.7% while reducing oxygen cost by only 3.2%. This reported gap between physiological and performance gains suggests that non-fatigued running economy assessments may capture only part of what AFT does to performance. This translation issue may have particular implications for women. Because the running economy-to-velocity relationship is speed-dependent (Kipp et al., 2019), sex-equivalent gains in running economy may yield larger race performance improvements in women, who compete at lower absolute speeds. It is therefore entirely plausible that men and women experience similar acute metabolic gains from AFT (Seglina et al., 2026), while the competitive value of those gains differs meaningfully between the sexes.

A second pathway is that women may derive genuinely greater physiological benefit from AFT, but only under conditions that short, fresh-state, fixed-intensity testing does not typically replicate. Fatigue is the most obvious candidate. Race performance depends on physiological responses under dynamic and progressively fatiguing conditions, and the physiological determinants of endurance performance, including running economy, deteriorate during prolonged exercise as athletes are challenged by fatigue (Hunter et al., 2025; Maunder et al., 2021; Jones, 2024). There is growing evidence that women may be relatively more fatigue-resistant in relevant endurance contexts (Tiller & Illidi, 2024; Besson et al., 2021). If AFT interacts with fatigue-related drift, neuromuscular deterioration, or the maintenance of propulsion later in exercise, then sex-equivalent running economy benefits observed in fresh-state testing may still translate to larger competitive gains in women. This may be especially relevant given that women complete a given distance in a longer absolute time than men, potentially increasing the opportunity for fatigue-sensitive footwear effects to emerge and amplify during competition. The principle that fresh-state testing can miss meaningful effects on running economy is not specific to footwear. Recent work demonstrated that 10 weeks of supplementary strength training produced no change in running economy when tested in a fresh state, yet reduced the deterioration in running economy over the final 30 minutes of a 90-minute run in well-trained male runners (Zanini et al., 2025). Interventions known to alter the determinants of endurance performance can therefore be entirely undetectable in standard laboratory protocols while exerting meaningful effects under fatigue.

Direct AFT-specific evidence of benefits under race-like fatigue remains limited and has not yet resolved whether these effects differ by sex. Schwalm et al. (2025) reported that AFT maintained its advantage over time without a progressive increase in benefit in a sample of nine runners. In contrast, Steele et al. (2025) observed that AFT attenuates neuromuscular fatigue and slows oxygen-uptake drift during prolonged running, and Black et al. (2022) reported that highly cushioned footwear reduced oxygen cost to a greater extent under fatigued conditions (∼4.7%) than in a fresh state (∼3.2%). Preliminary evidence from sprinting similarly suggests that AFT is effective when athletes must also maintain velocity against fatigue rather than simply attain it over shorter sprint distances (Mason et al., 2025). Together, these findings provide early support for the possibility that AFT benefits may be most pronounced when athletes are physiologically challenged; conditions that define road racing but remain largely absent from the current standard laboratory protocols.

The way testing intensity is matched between sexes may further influence whether sex-specific effects are detected. Because responses to new footwear technology vary with speed (Day & Hahn, 2020), the intensity-matching approach adopted may shape the results. Matties and Rowley (2025) reported that female collegiate runners showed a significantly greater metabolic benefit from AFT than males when self-selected race-pace running speed was controlled as a covariate, with no sex differences in the unadjusted analysis. In contrast, Seglina et al. (2026) found no sex difference when testing at matched relative intensities of 60-80% VO2peak. These divergent findings suggest that how intensity is handled, whether matched by physiological anchor, self-selected pace, or statistical adjustment, may influence whether sex-specific effects emerge. A broader pattern reinforces this gap: a recent meta-analysis found that only 4 of 17 AFT studies adjusted speeds by sex, and just 2 used time trials to assess actual running performance (Xiao et al., 2025). Future studies should therefore prioritise individualised, race-relevant intensities (Kuzmeski et al., 2026), and performance-based outcomes when assessing sex differences in AFT benefits.

Underpinning these issues is a more fundamental constraint: the continued underrepresentation of women in AFT research. As we recently reported, fewer than 15% of participants in published AFT studies are women (Mason et al., 2024). This pronounced imbalance leaves sex differences in AFT responses largely unresolved and increases the likelihood that meaningful female-specific effects may remain undetected in current experimental designs. Ultimately, most existing studies have simply been underpowered to assess sex-specific responses, even when women were included in the sample.

Overall, metabolic assessments allow highly controlled comparisons between footwear variations, offering a precise and replicable window into the key mechanism through which AFT contributes to performance. However, current evidence cannot resolve whether the larger competitive gains recently observed in women arise because women derive greater physiological benefit from AFT, because sex-similar metabolic gains translate more effectively to race-day performance for women, or because both processes occur together. Resolving this will require researchers to move beyond the convenience of short, fresh-state laboratory protocols and embrace the ecological complexity of real-world running. Multi-day, adequately powered trials to offset inter-day variability are unlikely to be convenient, but they may be pivotal in detecting the potential sex-specific effects that observational data, including our results, consistently suggest are there.

### Potential contributing factors beyond footwear

Although our findings support the interpretation that women have benefited more from AFT than men, we cannot exclude the possibility that other running-specific factors also contributed to the larger recent performance shifts observed in women’s road running. Importantly, however, these alternative explanations would need to account not only for the greater improvement in road running than throwing, but also for the absence of a comparable sex-specific pattern in the throwing events.

The most prominent such explanation is the broader, and overdue, expansion of women’s sport. The women’s Olympic marathon was first contested in only 1984, highlighting that women were permitted to enter elite endurance competitions far later than men, and were exposed to professionalisation, expanding competitive opportunity and rising participation over a compressed historical window (Cheuvront et al., 2005; Hallam & Amorim, 2022). There is direct evidence that such shifts narrow the sex gap in performance, with the ratio of male-to-female participants reportedly associated with the sex difference in marathon running velocity (Hunter & Stevens, 2013). In collegiate rowing, Title IX selectively incentivised women but not men, female participation rose without a corresponding change in men, accompanied by a narrowing of the performance gap (Keenan et al., 2018). We do not discount the continuing influence of these forces, which have recently been invoked to explain women’s ongoing performance gains independently of footwear (Jucker, 2026).

Our design speaks directly to this opportunity-legacy hypothesis. If broader improvement in women’s sport were the primary driver, its effects should not be confined to footwear-exposed events. A rising tide of this kind would be expected to lift women’s performances across the athletics programme. Yet we observed no comparable sex-divergent improvement in the throwing disciplines, where women’s gains matched men’s. If anything, this maturation logic runs counter to the data: the women’s hammer throw was contested at the Olympic Games only from 2000, more recently than the women’s marathon (1984), so an opportunity-driven account predicting continued catch-up in newer women’s events would anticipate strong gains in the women’s hammer, yet it was among the throwing events showing no significant improvement between eras. A further consideration concerns how such trends are measured: continuous annual mean-pace trajectories can obscure a rapid change in the composition of the elite field, and our churn analysis was designed to detect exactly this. Pre-AFT road-running performances have been almost entirely displaced from contemporary all-time lists, particularly in women’s events, whereas throwing events retain a substantial proportion of pre-AFT marks. This near-total, event-specific turnover is difficult to reconcile with the gradual, sport-wide trajectory an opportunity-legacy explanation implies, and instead points to a rapid shift concentrated in exactly the events where footwear changed.

Specific regional developments in elite women’s distance running are an accompanying explanation beyond footwear. Approximately 89% of the female road runners in our dataset originated from East Africa, whereas fewer than 10% of the throwing athletes in the current dataset hail from the same region. However any region-specific developments would need to have accelerated specifically from 2016 onwards, which is difficult to reconcile with the much longer-standing dominance of East African women in global distance running. The magnitude of pre-AFT performance displacement from current top 50 rankings further suggests an abrupt rather than gradual inflection, which is inconsistent with the longer timescales on which such regional developments typically operate.

Several further candidates warrant brief consideration, though none map cleanly onto the specific pattern we observe. Anti-doping detection, compliance, and effectiveness may differ by sex and region in ways that contribute to recent trends (Mason et al., 2024), and may be particularly important given the low proportion of adverse analytical findings relative to survey-based estimates of doping prevalence (WADA, 2024; Ulrich et al., 2018) and the recent region-specific controversies in road racing. Co-operative drafting strategies (Hoogkamer, Snyder & Arellano, 2018) may also disproportionately advantage women in elite road racing, though women have also improved more than men in sprint and middle-distance events (Mason et al., 2023; Willwacher et al., 2024) in the AFT era with only sex-equivalent pacing advantages available through the introduction of the wavelight during middle– and long-distance races. More speculative contributors, such as sex-specific responses to supplementation (Durkalec-Michalski et al., 2025), may also play some role, though women remain underrepresented in this literature too, including highly relevant sodium bicarbonate studies (Saunders et al., 2022).

Performance improvements between eras were not exclusive to road running. Several throwing events improved significantly, and the men’s shot put independently improved more than some men’s road-running disciplines despite no technological shift. Technique change is the most plausible explanation: in 2003, 7 of 10 World Championship finalists used the glide technique; by 2019, that figure was zero (Salinero & Del Coso, 2021), with the rotational technique’s takeover accelerating around 2016 (Hatase & Takanashi, 2022). The emergence of a generational athlete in Ryan Crouser may have contributed. Although the world record holder could only contribute one performance to our AFT era data pool, his performances may have elevated the entire field due to competitive pressure, in ways that have no parallel in women’s shot put over the same period. That women’s shot put has not followed the same trajectory likely reflects a slower uptake of the rotational technique and a younger average age among female rotation throwers (Hatase, Okada & Takanashi, 2025), suggesting the performance gains may be delayed rather than absent. The finding that our control was itself sensitive to broader athletic trends such as these, rather than remaining passive, confirms that throwing events were responsive to the concurrent forces shaping elite athletics, making the markedly larger and sex-specific gains in road running even more striking by comparison.

### Limitations

Several limitations should be considered when interpreting these findings. First, our analysis was restricted to the top 50 performances per event and era, meaning the conclusions are specific to elite-level competition and may not generalise to broader performance populations. Second, while throwing events offered a useful non-AFT control family, they remain an imperfect comparator, and event-specific developments such as the shot-put technique revolution can affect throwing in ways that are neither footwear-related nor captured by the broad athletic trends we sought to control for. Third, the use of World Athletics points provided a practical means of standardising performance across disciplines, but this approach does not fully eliminate the complexity of comparing fundamentally different event types. Moreover, our study period included two Olympic Games in the AFT era compared to only one Olympic Games in the pre-AFT era. Given that global performances typically show an uptick during Olympic years due to increased competition and athlete peaking strategies (Haake, Foster & James, 2013), this imbalance may contribute to the observed performance improvements independent of footwear technology effects. Finally, although our design improves on prior observational analyses by incorporating a control family of events, it remains observational in nature and cannot fully isolate the independent contribution of footwear technology from all other potential influences.

## Conclusion

We provide the most controlled and compelling observational evidence yet of sex-specific performance responses to AFT, finding that performance gains in the AFT era are larger in road running than in throwing, and are disproportionately greater in women than in men. While factors beyond footwear may contribute to recent endurance performance evolution, the absence of comparable sex-divergent trends in throwing events suggests that broader improvements in women’s sport are unlikely to fully explain the magnitude and specificity of the observed recent sex gap in road running. The near-total displacement of pre-AFT performances from contemporary road-running all-time lists reinforces this interpretation, pointing to a rapid and event-specific shift that is difficult to reconcile with more gradual shifts in women’s sport.

These results strengthen the case that sex-specific responses to AFT warrant further experimental investigation, particularly given that the vast majority of experimental studies have not been purposely designed to detect sex-specific responses, and that women remain systematically underrepresented in AFT research. Further advances will require larger female sample sizes, elegant approaches to intensity and metabolic matching, and experimental designs that better reflect the demands of competitive running, including fatiguing and race-relevant protocols, and performance-based outcomes in field settings. Ultimately, reconciling experimental and ecological evidence will be essential for understanding how, and for whom, AFT confers its greatest competitive advantage.

## Author Contributions

**Conceived research:** JM

**Designed research:** JM, LB

**Analysed data:** LB, DH

**Interpreted results of analysis:** JM, LB, DH

**Prepared figures:** LB

**Drafted manuscript:** LB, DH, RT, LH, JM

**Edited and revised manuscript:** LB, DH, RT, LH, JM

**Approved final version:** LB, DH, RT, LH, JM

**Supervision:** JM

## Acknowledgements

The authors would like to thank Dr Sam Cheuvront for his insightful feedback and discussions in the end stages of this manuscript, and the Trackademic junior research team for their spirited feedback throughout the research process.

